# Genomic Evidence for a-α Heterothallic and α-α Unisexual Mating and Recombination in an Environmental *Cryptococcus deneoformans* Population

**DOI:** 10.1101/2025.08.20.671223

**Authors:** Megan Hitchcock, Veronica Thorn, Himeshi Samarasinghe, Sheng Sun, Joseph Heitman, Jianping Xu

**Affiliations:** Department of Biology, McMaster University, Hamilton, ON L8S 4K1, Canada; Department of Molecular Genetics & Microbiology, Duke University, Durham, NC 27710, United States

**Keywords:** Cryptococcosis, single nucleotide polymorphisms, phylogenetic incompatibility, four-gamete test, mating type locus, mitochondrial recombination

## Abstract

*Cryptococcus deneoformans* is a human fungal pathogen capable of both **a**-α and α-α mating and sexual reproduction in laboratory settings. However, the extent of **a**-α and α-α sexual reproductions in natural populations remain unexplored. Here we analyzed the whole-genome sequences of 24 environmental strains of *C. deneoformans* from western Saudi Arabia, including one *MAT***a** and 23 *MAT*α isolates, with 15 *MAT*α isolates belonging to multi-locus sequence type ST160 as defined by their combined DNA sequences at seven loci. To identify signatures for **a**-α and α-α reproduction, three samples were analyzed: total, *MAT*α, and ST160. For each subpopulation, single nucleotide polymorphisms (SNPs) were identified for both the nuclear and mitochondrial genomes and subjected to four-gamete tests. In the total population and the *MAT*α subpopulation, variable proportions of SNP pairs within as well as between the nuclear and the mitochondrial genomes showed evidence for recombination. Though no mitogenome SNPs were found among the 15 strains of ST160, the nuclear genome showed clear evidence for recombination, including among SNPs within the mating type region. In addition, the nuclear genome SNP pairs located further apart on the same chromosome showed a greater frequency of recombination in all three sample types. In contrast, mitogenome recombination breakpoints were mainly located in two genomic regions. Together, these results provide robust evidence for both **a**-α and α-α sexual reproduction within this environmental population of *C. deneoformans*.

**Article Summary:** This study compares conclusions drawn from multilocus sequence data and whole-genome SNP data regarding genetic diversity and reproduction in an environmental population of *Cryptococcus deneoformans*. By analyzing bi-allelic SNPs across 24 strains isolated from western Saudi Arabian soils, evidence of both **a**-α and α-α recombination was found, suggesting the occurrence of both **a**-α and α-α sexual reproduction in nature. This provides strong support for these reproductive modes and indicates diverse mechanisms of genetic exchange in fungal pathogens in nature. Additionally, our results suggest that multi-locus sequence typing underestimates genetic diversity, potentially confounding interpretations of population structure and evolutionary history.

## Introduction

The *Cryptococcus neoformans* species complex (*CNSC*) is a globally distributed group of pathogenic basidiomycetous yeasts that were recently classified by the World Health Organization (WHO) as a critical priority fungal pathogen [1–5]. *CNSC* causes cryptococcosis through the inhalation of spores or desiccated vegetative yeast cells from environmental sources such as avian excreta, soil, and trees [6–8]. Each year globally, about 200,000 individuals suffer from cryptococcal meningitis, causing about 147,000 deaths [9]. Based on differences in cell surface antigens and genetic factors, *CNSC* is divided into two divergent serotypes, A and D, corresponding to two species, *C. neoformans* and *C. deneoformans*, respectively, and their hybrids, serotype AD [10–12].

Although *C. deneoformans* causes <5% of global cryptococcosis, it can be highly prevalent in specific regions, such as around the Mediterranean [13]. Despite a genetic divergence of up to 15% at the whole-genome level, *C. neoformans* and *C. deneoformans* can undergo hybridization, generating serotype AD interspecies hybrids [14–16]. Recently, Cogliati et al. analyzed a global collection of 102 serotype AD hybrids from across five continents [17]. This analysis revealed evidence for both ancient and recent hybridization events between serotypes A and D strains in many regions and suggests the distribution of *C. deneoformans* could be much broader than previously thought [17].

*C. deneoformans* has a well-defined bipolar mating system but has been hypothesized to reproduce predominantly clonally in nature, through either asexual budding or unisexual selfing. This notion is supported by the overrepresentation of a few multilocus genotypes across many regions and that most loci in *C. deneoformans* populations seemed in linkage disequilibrium [18,19]. Mating in *C. deneoformans* is determined by the mating-type locus (*MAT)*, which has two distinct mating types (**a** and α), with no mating type switching [18]. Under mating-inducing conditions, **a**-α mating typically generates equal frequencies of *MAT***a** and *MAT*α progeny [20–22] and demonstrates uniparental mitochondrial DNA (mtDNA) inheritance, with progeny primarily inheriting mtDNA from the *MAT***a** parent [23]. In contrast, during α-α unisexual reproduction, while the recombination frequency in the nuclear genome is reduced by ∼2-fold compared to **a**-α sexual reproduction [24,25], the mitochondrial genome (mitogenome) is biparentally inherited with evidence of recombination. For unisexual reproduction, α-α mating happens more readily than the **a-a** mating in the laboratory [26], which could have contributed to the skew in mating type distribution observed in nature in favor of *MAT*α strains. A previous study of the sister species *C. neoformans* found that the decay of linkage disequilibrium occurs in *C. neoformans* lineages with a higher prevalence of both mating types (VNBI and VNBII) as well as those with few if any *MAT***a** isolates (VNI), providing population genomic evidence for both modes of sexual reproduction in nature [27]. However, the extent of **a**-α and α-α sexual reproduction in natural populations of *C. deneoformans* remains largely unexplored.

Samarasinghe et al. analyzed 562 soil samples collected from six sites across Saudi Arabia and obtained 76 isolates of *C. deneoformans* at two sites (Jeddah and Yanbu) along the Red Sea in western Saudi Arabia [28]. Based on DNA sequences at seven loci used by the *Cryptococcus* research community as the consensus multi-locus sequence typing [29], Samarasinghe et al. identified this population as predominantly clonal, with limited evidence for recombination, and described the population as the first desert environmental reservoir of *C. deneoformans* isolates [28].

Because the seven loci used in the consensus MLST scheme are highly conserved housekeeping genes, their sequence variations likely underrepresent the whole-genome genetic diversity. In this study, we examined the whole genome sequences among 24 Saudi Arabian isolates to further explore the relationships among single nucleotide polymorphisms. These 24 isolates represent all six MLST genotypes, both geographic regions, and the phenotypic diversities as reported for the original 76 Saudi Arabian isolates. Our population genomic analyses identified large cryptic genetic variations, including evidence for both **a**-α and α-α sexual reproduction within this environmental population.

## Materials and Methods

### Strains included in the study and whole genome sequencing

Twenty-four strains of *C. deneoformans* were analyzed in this study. These isolates were collected from two western regions of Saudi Arabia: Jeddah (6 strains) and Yanbu (18 strains) as summarized in **Table 1**. Most isolates (23/24; 95.8%) were mating type *MAT*α with a single isolate being *MAT***a**. The multilocus genotypes of these isolates were previously identified based on a consensus multi-locus sequence typing scheme, which assigns a genotype based on the combined sequences at seven loci [29,30]. Among the 24 isolates, 15 belonged to the same MLST genotype ST160, five were identified as ST294, and the remaining four were individually identified as ST449, ST613, ST614, and ST615 in the cryptococcal MLST database (**Table 1**). While ST160, ST294 and ST449 have been reported from other geographic regions, ST613, ST614, and ST615 have only been reported from Saudi Arabia [31].

**Table 1.** List of isolates analyzed in this study. Their sampling location, MLST genotype, mating type, and average sequencing depth for each strain after filtering. The single *MAT***a** strain is bolded.

| Strain ID | Sampling Location | MLST ST No. | Mating Type | Average Sequence Depth |
| --- | --- | --- | --- | --- |
| J1 | Jeddah | 160 | $\alpha$ | 81 |
| J4 | Jeddah | 160 | $\alpha$ | 95 |
| J7 | Jeddah | 160 | $\alpha$ | 58 |
| J20 | Jeddah | 160 | $\alpha$ | 89 |
| J26 | Jeddah | 160 | $\alpha$ | 86 |
| J86 | Jeddah | 160 | $\alpha$ | 80 |
| Y30 | Yanbu | 160 | $\alpha$ | 89 |
| **Y38** | **Yanbu** | **449** | **a** | **87** |
| Y87 | Yanbu | 615 | $\alpha$ | 83 |
| Y97 | Yanbu | 614 | $\alpha$ | 83 |
| Y102 | Yanbu | 160 | $\alpha$ | 107 |
| Y106 | Yanbu | 160 | $\alpha$ | 100 |
| Y107 | Yanbu | 294 | $\alpha$ | 88 |
| Y108 | Yanbu | 294 | $\alpha$ | 89 |
| Y111 | Yanbu | 160 | $\alpha$ | 90 |
| Y112 | Yanbu | 613 | $\alpha$ | 93 |
| Y113 | Yanbu | 160 | $\alpha$ | 98 |
| Y122 | Yanbu | 160 | $\alpha$ | 86 |
| Y132 | Yanbu | 160 | $\alpha$ | 103 |
| Y157 | Yanbu | 294 | $\alpha$ | 74 |
| Y163 | Yanbu | 160 | $\alpha$ | 109 |
| Y164 | Yanbu | 294 | $\alpha$ | 90 |
| Y184 | Yanbu | 160 | $\alpha$ | 97 |
| Y187 | Yanbu | 294 | $\alpha$ | 82 |

To obtain their whole-genome sequences, the 24 isolates were first sub-cultured on Yeast Extract-Peptone-Dextrose agar (YEPD) and grown for two days at 30°C. DNA was extracted following the protocol described by [31]. Extracted genomic DNA was sent for sequencing at Duke University’s genomics center. The average read sequence depth ranged from 58-107 folds of the genome for the 24 strains. The raw sequence reads for the 24 strains have been deposited in GenBank under BioProject number PRJNA1298968 (https://www.ncbi.nlm.nih.gov/bioproject/PRJNA1298968), with sequence read archive accession numbers SRR34787920 to SRR34787943.

### Pre-processing and Variant Calling

The raw sequence data for each isolate was checked for quality and trimmed with Fastp (v0.23.1) [30]. Reads were kept only when at least 60% of bases had a phred score of 30 or higher and with a minimum length of 30 base pairs. The filtered reads were aligned to the reference genome of *C. deneoformans* strain JEC21 (GCA_000091045.1) with the Burrows-Wheeler Aligner (BWA-MEM v0.7.17-r1188) [32]. The alignments were further processed with samtools (v1.17) [33] to exclude sequences with low mapping quality and length. Duplicate reads were removed with MarkDuplicates in the Picard tool and genetic variants were called across the whole genome with the GATK tool suite, including HaplotypeCaller (with the ‘-ERC GVCF’ flag), GenomeDBImport, and GenotypeGVCFs [34]. The resultant multisample VCF file was subjected to additional filtering with bcftools [33] to remove non-biallelic SNP loci, loci that fell within known repetitive regions and centromeres of the genome. For recombination tests, only SNP loci with a minor allele count in at least two isolates in our population sample were kept. Aside from the total sample of 24 isolates for analyses, two additional subpopulations were extracted and analyzed separately: one subpopulation included only the 23 *MAT*α isolates, and the second subpopulation included only the 15 isolates previously identified as belonging to ST160 based on multi-locus sequence typing. The mitochondrial dataset was performed similarly, where the reads were aligned to an assembled mitochondrial reference genome of JEC21 [35].

### Phylogenetic Analyses

Phylogenetic analyses were performed on the concatenated SNPs of the nuclear and mitochondrial genomes, respectively, to investigate the relationship among strains. Additionally, a phylogenetic analysis was performed on the concatenated SNPs located within the *MAT* locus. In all phylogenetic analyses, SNPs were aligned with ClustalW to construct a maximum parsimony tree through PAUP* [36–38]. All trees were rooted with the *C. deneoformans* reference strain JEC21 and visualized with an interactive tree of life (iTOL) program [39]. The consistency index and homoplasy index of the nuclear, mitochondrial, and *MAT* trees were calculated with the PAUP* software [38].

### Four Gamete Test

To evaluate the degree of recombination in the three population samples, we calculated the percentages of SNP pairs that were phylogenetically incompatible. Phylogenetically incompatible SNP pairs in populations of organisms are commonly used as evidence of recombination. Phylogenetic incompatibility was assessed with the four gametes test. The test was performed between all filtered bi-allelic SNP pairs in the total population and in the two subpopulations. For each test, a binary SNP matrix was generated with the bcftools query on the filtered VCF file. The test was performed with a custom C program (available at https://github.com/thorn-v/FGT) resulting in a large list of SNP pairs which display all four gamete combinations in the given population. This file was then processed to extract relevant SNP pairs (e.g., intrachromosomal pairs) utilizing standard Unix tools. Subsequent analysis and visualization were performed in R with the ggplot2 [40] and circlize [41] packages. Difference in the frequencies of intra-chromosomal and inter-chromosomal SNP pairs being phylogenetically incompatible were tested via Wilcoxon rank sum exact test [42].

We further tested the hypothesis that if recombination existed in various populations, the SNP pairs located further apart on each chromosome should recombine more often and thus show a higher frequency of phylogenetic incompatibility and recombination than those located closer together on the chromosome. The relationships between physical distance of SNP pairs within individual chromosomes and the frequency of phylogenetic incompatibility were assessed with R and visualized with ggplot2 [40]. Because the mitogenome is a circular molecule, the shorter of the two possible physical distances between SNPs on the mitogenome were used in our assessment of their relationships between physical distance and the frequency of recombination.

### Mating

To examine the fertility of isolates and if mating could be induced within laboratory conditions, all 24 strains were crossed with standard tester strains JEC20 (*MAT***a**) and JEC21 (*MAT*α). In preparation of mating, all strains were first cultured on Yeast Extract-Peptone-Dextrose (YEPD) agar medium for 2–3 days at 30 °C. A toothpick of actively growing cells was re-suspended in 100ul of sterile distilled water and 5 μl of both parental strains were sequentially layered onto V8-juice mating agar medium along with 10 μl of each strain individually to serve as negative controls. All strains were crossed with both JEC20 and JEC21 to test for **a**-α and α-α mating. V8-juice mating agar medium was prepared with 40 g agar, 50 μl V8 juice and 950 μl sterile distilled water and adjusted to pH=5. To allow for mating, these plates were incubated in the dark at room temperature (23 °C) and checked for evidence of mating (the formation of hyphae) after three weeks.

## Results

### WGS Reveals Extensive Sequence Variation across the Genomes of the Saudi Arabian Isolates

#### Nuclear genome SNPs

The read depth from whole-genome sequencing ranged from 58 to 109 folds among the 24 strains (**Table 1**). With the assembled nuclear genome sequence of *C. deneoformans* strain JEC21 as a reference (GCA_000091045.1), a total of 203,467 SNPs were identified among the 24 strains. Among these SNPs, 79,541 had the rare allele found in only one of the 24 strains, the remaining 104,793 SNPs had the rare allele found in at least two of the 24 strains (i.e. phylogenetically informative SNPs, PI-SNPs). Because the 79,541 SNPs with singleton alleles could have been derived from spontaneous mutations accumulated during culturing, and errors from sequencing library preparation, and/or during sequencing, our analyses will focus on the 104,793 PI-SNPs for the remainder of the analyses.

These 104,793 SNPs were distributed across all chromosomes and the number of SNPs varied among the chromosomes, ranging from 4,259 on chromosome 13 to 13,512 on chromosome 1 (**Table 2**), and their frequencies ranged from 0.4329% (on chromosome 10) to 0.6282% (on chromosome 11). Among the 23 *MAT*α isolates, 94,447 SNPs with the rare allele present in at least two strains were identified at the whole genome level, with counts ranging from 2,621 on chromosome 10 to 12,090 on chromosome 1 (**Table 2**), and their frequencies ranged from 0.2414% (on chromosome 10) to 0.5575% (on chromosome 11). Within the ST160 subpopulation, a total of 2,027 SNPs with the rare allele present in at least two strains were detected, varying from 26 SNPs on chromosome 12 to 302 SNPs on chromosome 2 (**Table 2**), with their frequencies ranging from 0.0029% (on chromosome 12) to 0.0203% (on chromosome 14). Within each chromosome, there is large variation among regions in their SNP densities in all three sample types (**Figure 1**). Two of the three sample types, the total population (N=24) and the *MAT*α population (N=23), showed very similar SNP density distributions across almost all chromosomes. Not surprisingly, the ST160 population (N=15) showed lower SNP density across all chromosomes and chromosomal segments than the other two populations. However, the major SNP density peaks in the ST160 population corresponded to most of the peaks found in the total and the *MAT*α populations (**Figure 1**).

**Figure 1.**
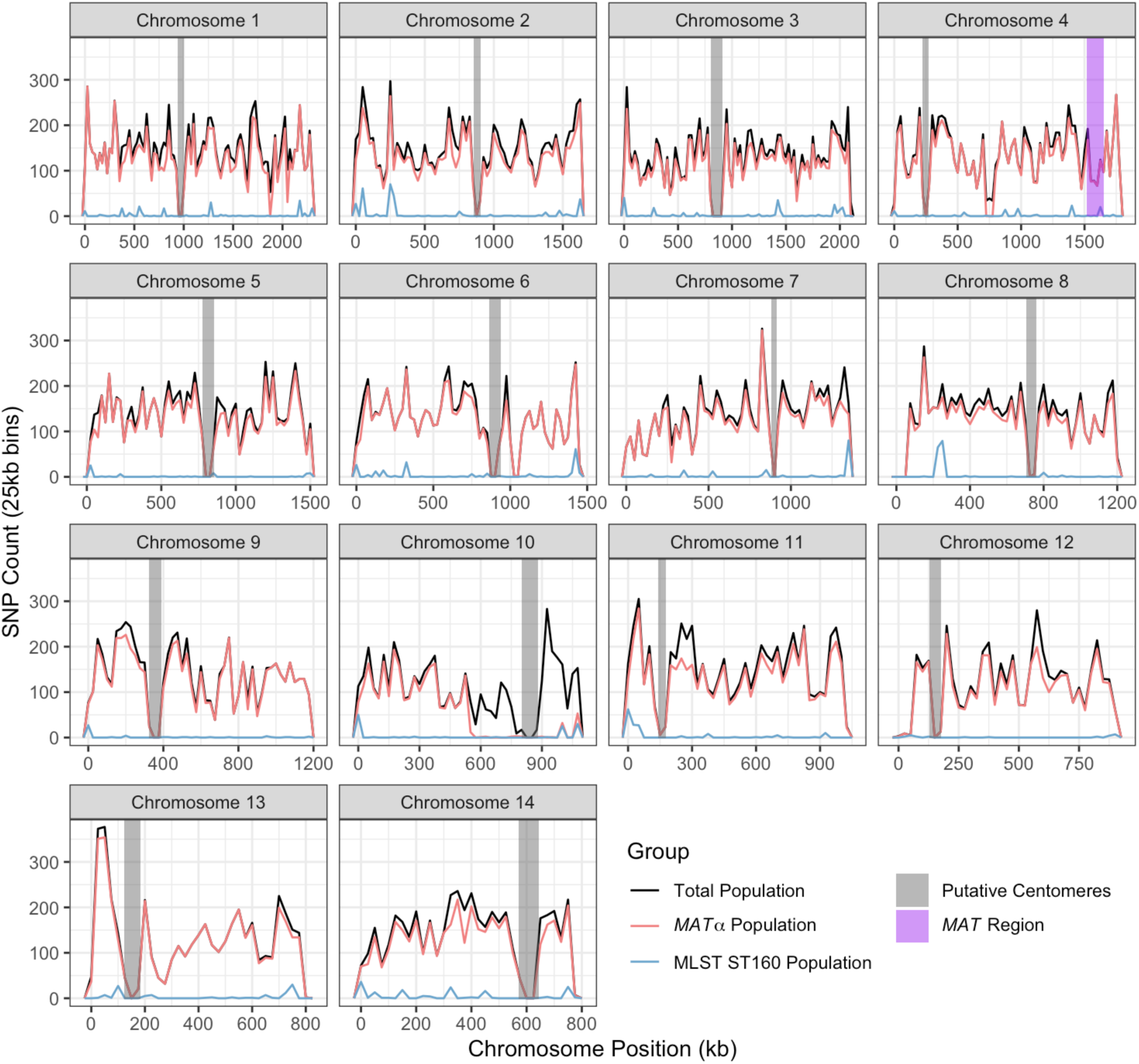
Distribution of single nucleotide polymorphisms (SNPs) among 24 Saudi Arabian strains across each chromosome. Chromosome position is shown in kilobases (kb) on the X-axis while SNP counts are displayed in 25 kilobase bins on the Y-axis on each chromosome. Three different populations are represented, the total population (N=24) in black, the *MAT*α population (N=23) in pink, and the ST160 population (N=15) in blue. Putative centromere regions are denoted in grey on each chromosome and the *MAT* region in purple on chromosome 4.

**Table 2.**
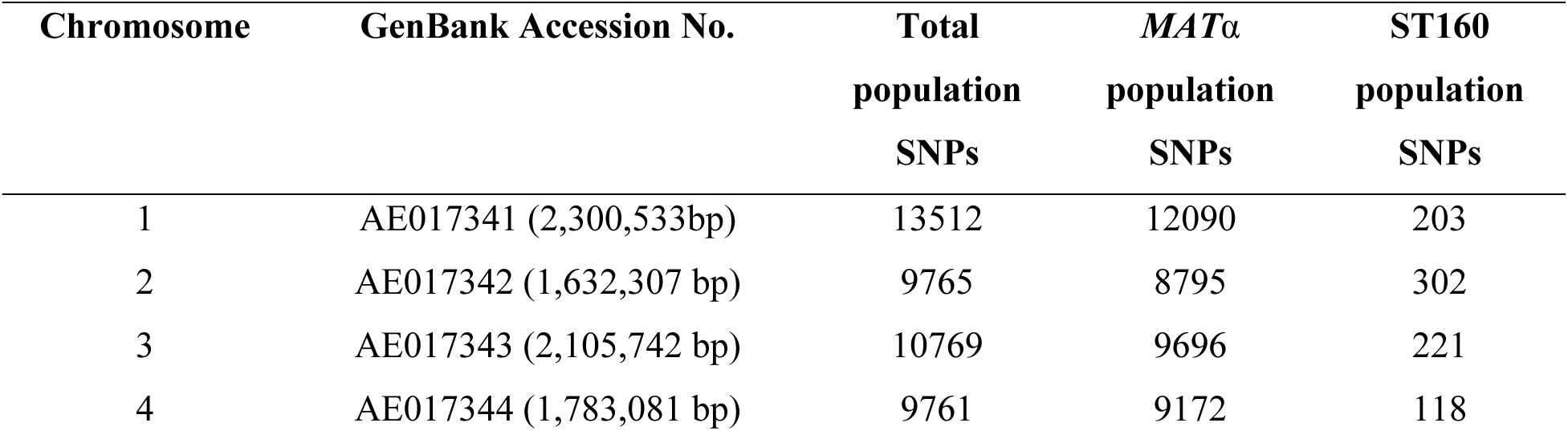
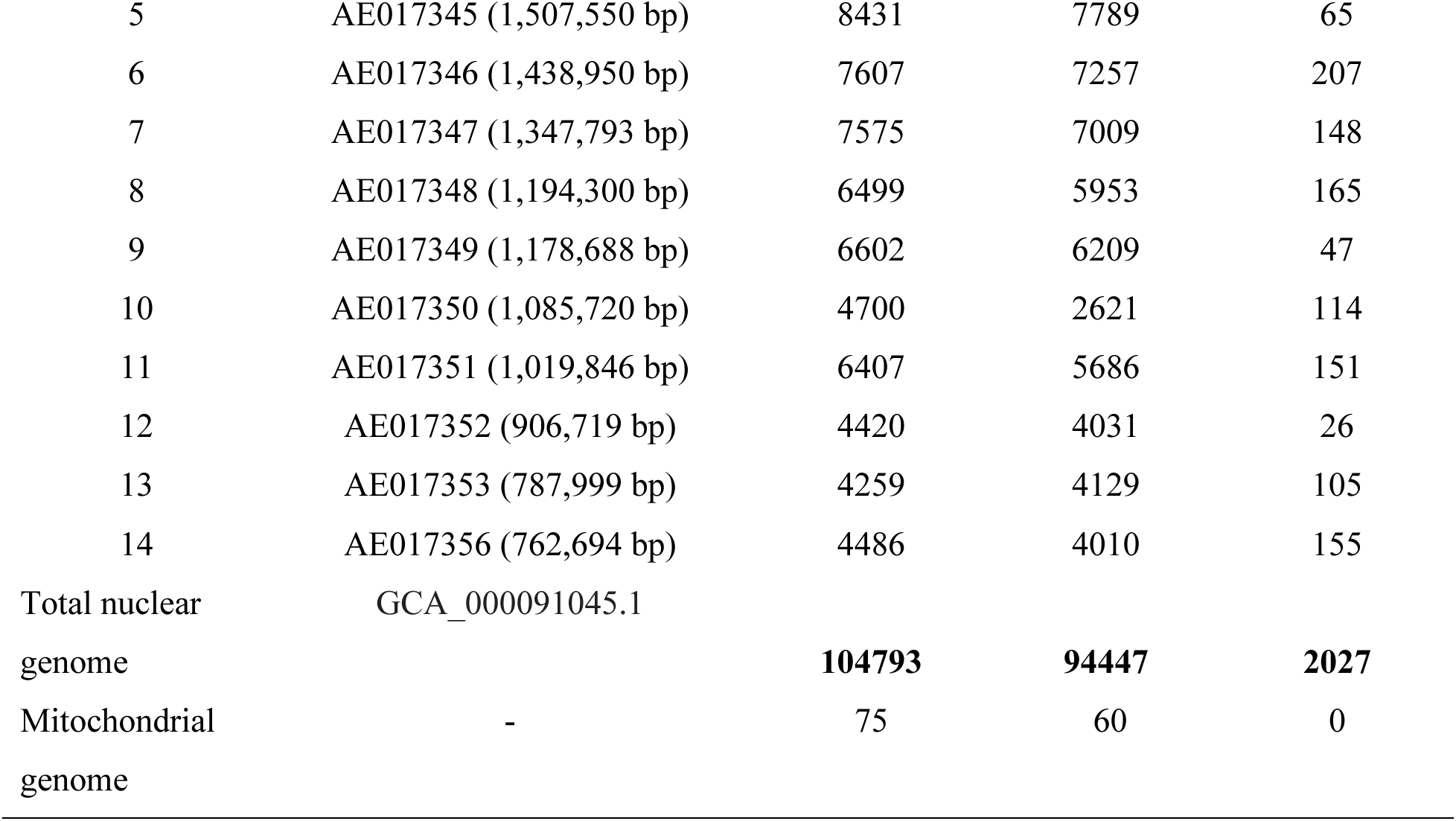
Summary numbers of filtered bi-allelic SNPs identified when compared to reference genome of strain JEC21. Only SNPs with the minor allele being in at least two isolates in each population are presented in this table and for the four-gamete tests. The nuclear chromosome number, GenBank accession number of each chromosome, and size of chromosome (in parenthesis) represented those of strain JEC21. The numbers of SNPs identified for the three population samples are shown in three columns, the total population, the *MAT*α and the ST160 subpopulations.

| Chromosome | GenBank Accession No. | Total<br>population<br>SNPs | *MATα*<br>population<br>SNPs | ST160<br>population<br>SNPs |
| --- | --- | --- | --- | --- |
| 1 | AE017341 (2,300,533bp) | 13512 | 12090 | 203 |
| 2 | AE017342 (1,632,307 bp) | 9765 | 8795 | 302 |
| 3 | AE017343 (2,105,742 bp) | 10769 | 9696 | 221 |
| 4 | AE017344 (1,783,081 bp) | 9761 | 9172 | 118 |
| 5 | AE017345 (1,507,550 bp) | 8431 | 7789 | 65 |
| 6 | AE017346 (1,438,950 bp) | 7607 | 7257 | 207 |
| 7 | AE017347 (1,347,793 bp) | 7575 | 7009 | 148 |
| 8 | AE017348 (1,194,300 bp) | 6499 | 5953 | 165 |
| 9 | AE017349 (1,178,688 bp) | 6602 | 6209 | 47 |
| 10 | AE017350 (1,085,720 bp) | 4700 | 2621 | 114 |
| 11 | AE017351 (1,019,846 bp) | 6407 | 5686 | 151 |
| 12 | AE017352 (906,719 bp) | 4420 | 4031 | 26 |
| 13 | AE017353 (787,999 bp) | 4259 | 4129 | 105 |
| 14 | AE017356 (762,694 bp) | 4486 | 4010 | 155 |
| Total nuclear<br>genome | GCA_000091045.1 | **104793** | **94447** | **2027** |
| Mitochondrial<br>genome | - | 75 | 60 | 0 |

Among the SNP density comparisons within and among chromosomes, the most notable difference was observed on the right arm of chromosome 10. Here, from ∼500,000 bp to ∼1,100,000 bp of chromosome 10, we observe a high density of SNPs in the total population but not in either of the two subpopulations (**Figure 1**). Further analyses revealed that 1,865 of the 1,898 SNPs located in this region for the total population are shared only between sample Y38, the *MAT***a** isolate, and isolate Y87 (*MAT*α). The high degree of regional sequence similarity between the *MAT***a** isolate Y38 and the *MAT*α isolate Y87 is consistent with recombination between *MAT***a** and *MAT*α isolates in this population.

#### Mitogenome SNPs

Similar to identifying SNPs in the nuclear genomes, we used the mitogenome of *C. deneoformans* JEC21 as reference for identifying mitogenome SNPs (mtSNPs) in our samples. Our analyses identified 159 total mtSNPs among the 24 isolates; 104 mtSNPs among the 23 *MAT*α isolates; and no mtSNP was found among the 15 ST160 isolates. Among these, there were 75 phylogenetically informative mtSNPs (i.e. PI-mtSNPs) in the total population and 60 PI-mtSNPs in the *MAT*α subpopulation of 23 isolates. In total, based on all the mtSNPs, nine mitochondrial genotypes were identified among the 24 isolates. All 15 ST160 strains shared one mitogenome type, while two of the five ST294 isolates (Y111 and Y157) also shared a mitogenome type. The remaining seven mitogenome types were each found in a single isolate.

### Sequence Analyses Provide Evidence for Sexual Reproduction

#### Phylogenetic evidence for recombination

The relationships among the isolates were analyzed based on the concatenated SNPs separately for the mitochondrial and nuclear genomes with a maximum parsimony method (**Figure 2**). The two phylogenetic trees revealed a similar clustering pattern, distinguishing the ST160 subpopulation from the remaining isolates and with isolate Y87 being the most distant from the other isolates. The 15 ST160 isolates were clustered together in both phylogenetic trees. These 15 isolates were from two geographic regions Yanbu and Jeddah, consistent with long distance dispersal of this genotype. Similarly, the five ST294 isolates were clustered together. Interestingly, for the mitochondrial phylogeny, isolate Y112 (ST613) was clustered with the five ST294 isolates. For the nuclear phylogeny, two isolates, Y112 (ST613) and Y97 (ST614), were clustered with the five ST294 isolates. The results are consistent with different ancestries of the nuclear and mitochondrial genes and genomes for different isolates.

**Figure 2.**
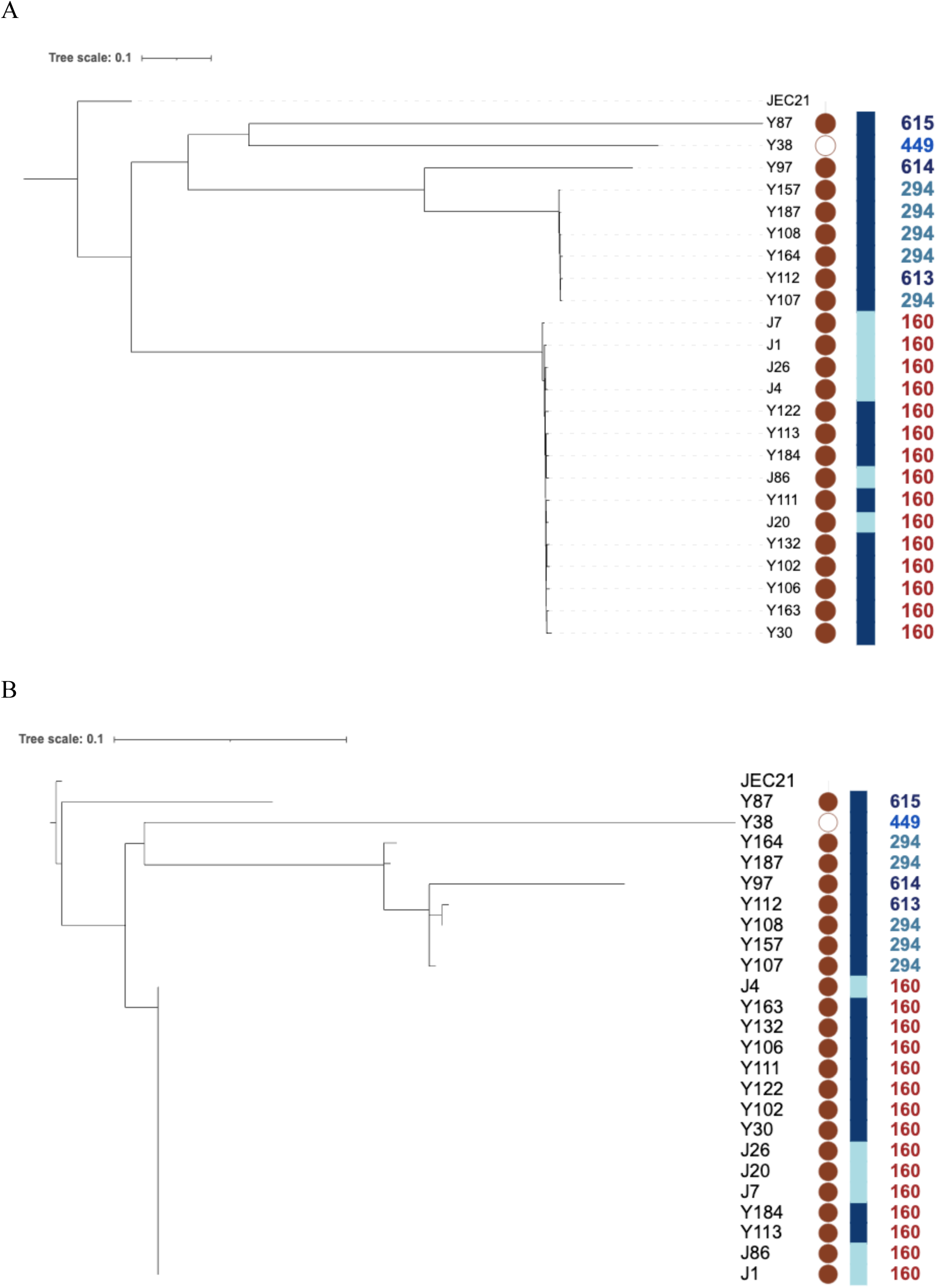
Phylogenetic analyses of 24 strains of *C. deneoformans* from Saudi Arabia. **A.** Phylogenetic analysis of nuclear SNPs conducted with maximum parsimony method. **B.** Phylogenetic analysis of mitochondrial SNPs analyzed using the maximum parsimony method. Within each tree, the symbols in columns left to right represent: (i) mating-type locus (*MAT*) where filled circles represent *MAT*α and empty circle represents *MAT***a**; (ii) sampling location (SL): Jeddah is represented by pale blue and Yanbu is represented by navy blue; and (iii) MLST sequence type numbers: ST160 isolates represented in brown, ST294 in green, ST449 in royal blue and the remaining three genotypes represented by dark blue.

The consistency index (CI) for both the nuclear (0.734) and mitochondrial (0.809) maximum parsimony trees deviated from 1. CI is measured by dividing the fewest possible evolutionary steps required to remain consistent with the data by steps taken to reconstruct the most likely observed relationship. In a completely clonal population while assuming no parallel mutation or reversion, the consistency index should equal to 1. Without parallel mutation or reversion, a CI less than 1 would be indicative of recombination, with lower CI values corresponding to greater evidence and higher frequency of recombination Our observed CI values here are consistent with recombination playing an important role in both the nuclear and the mitochondrial genomes in this environmental population.

#### Four-gamete test of nuclear SNP pairs

To investigate recombination patterns within this population, phylogenetic incompatibility via four-gamete tests was assessed for various population samples. For the nuclear genome, this analysis was conducted across all inter- and intra-chromosomal SNP pairs. The results are summarized in **Table 3**. In the total population (N=24), 19.38% of the genome-wide SNP pairs were found to be phylogenetically incompatible, while intrachromosomal SNP pairs ranged from 7.90% (chromosome 11) to 21.45% (chromosome 5). In the *MAT*α population (N=23; i.e., excluding the *MAT***a** isolate Y38, of ST449), removal of the *MAT***a** isolate and re-filtering to retain SNPs with minor allele shared by at least two strains of the *MAT*α population resulted in ∼10% decrease in total SNPs analyzed.

**Table 3.** Summary of the proportions of intra-chromosomal SNP pairs that are phylogenetically incompatible as determined by the four-gamete test. The proportions were determined for each chromosome in each of three populations: the total population, the *MAT*α, and the ST160 subpopulations. Chromosome and mitochondrial genome percentages represent intra-chromosomal SNP pairs that are phylogenetically incompatible. Total nuclear genome SNP pairs being phylogenetically incompatible includes both intra- and inter-chromosomal SNP pairs.

| Chromosome | Full Population SNP<br>pairs in PI | *MATα* SNP pairs<br>in PI | ST160 SNP pairs in<br>PI |
| --- | --- | --- | --- |
| 1 | 20.31% | 8.84% | 25.20% |
| 2 | 13.72% | 6.40% | 24.12% |
| 3 | 12.88% | 5.99% | 19.29% |
| 4 | 20.89% | 8.08% | 16.53% |
| 5 | 21.45% | 9.82% | 13.75% |
| 6 | 18.16% | 8.13% | 30.18% |
| 7 | 13.27% | 4.55% | 20.92% |
| 8 | 13.46% | 5.26% | 44.55% |
| 9 | 17.32% | 8.59% | 11.38% |
| 10 | 12.10% | 2.96% | 8.97% |
| 11 | 7.90% | 1.39% | 8.46% |
| 12 | 19.85% | 10.72% | 27.38% |
| 13 | 15.40% | 8.91% | 21.06% |
| 14 | 18.75% | 9.22% | 38.79% |
| Total Nuclear Genome | 19.38% | 8.76% | 23.49% |
| Mitochondrial Genome | 36.3% | 22.3% | NA |

Interestingly, such an exclusion led to a notable reduction in genome-wide phylogenetically incompatible SNP pairs, reducing the total percentage to 8.76%, about half of that in the total population. The percentages of intrachromosomal pairs that were phylogenetically incompatible were similarly reduced by about half compared to the full population of 24 isolates.

To further explore recombination patterns within this population, we tested for signatures of recombination within the ST160 subpopulation. Here, the four-gamete test was performed on all SNP pairs with the rare allele of each SNP locus present in at least two of the 15 isolates. This analysis revealed a very high level of phylogenetic incompatibility, with 23.49% of genome-wide nuclear SNP pairs being phylogenetically incompatible. The intrachromosomal SNP pairs showed incompatibility rates ranging from 8.46% to 44.55% (**Table 3**). Together, the results indicate signatures of recombination are present within the ST160 *MAT*α subpopulation.

The proportions of SNP pairs displaying all four gamete combinations are illustrated in **Figure 3**. In all three populations, not surprisingly, there was an overall higher proportion of SNP pairs on different chromosomes being phylogenetically incompatible than those on the same chromosomes. However, the difference was statistically significant only in the total population (p < 0.05, **Figure 3A**). As shown in **Figure 3B**, phylogenetic incompatibility was observed for SNP pairs across all chromosomes. Among the three populations, the highest proportion of SNP pairs showing phylogenetic incompatibility was in the ST160 population, which was followed by the total population, and the *MAT*α population, for both the overall intra-chromosomal SNP pairs and the inter-chromosomal SNP pairs (**Figure 3B**).

**Figure 3.**
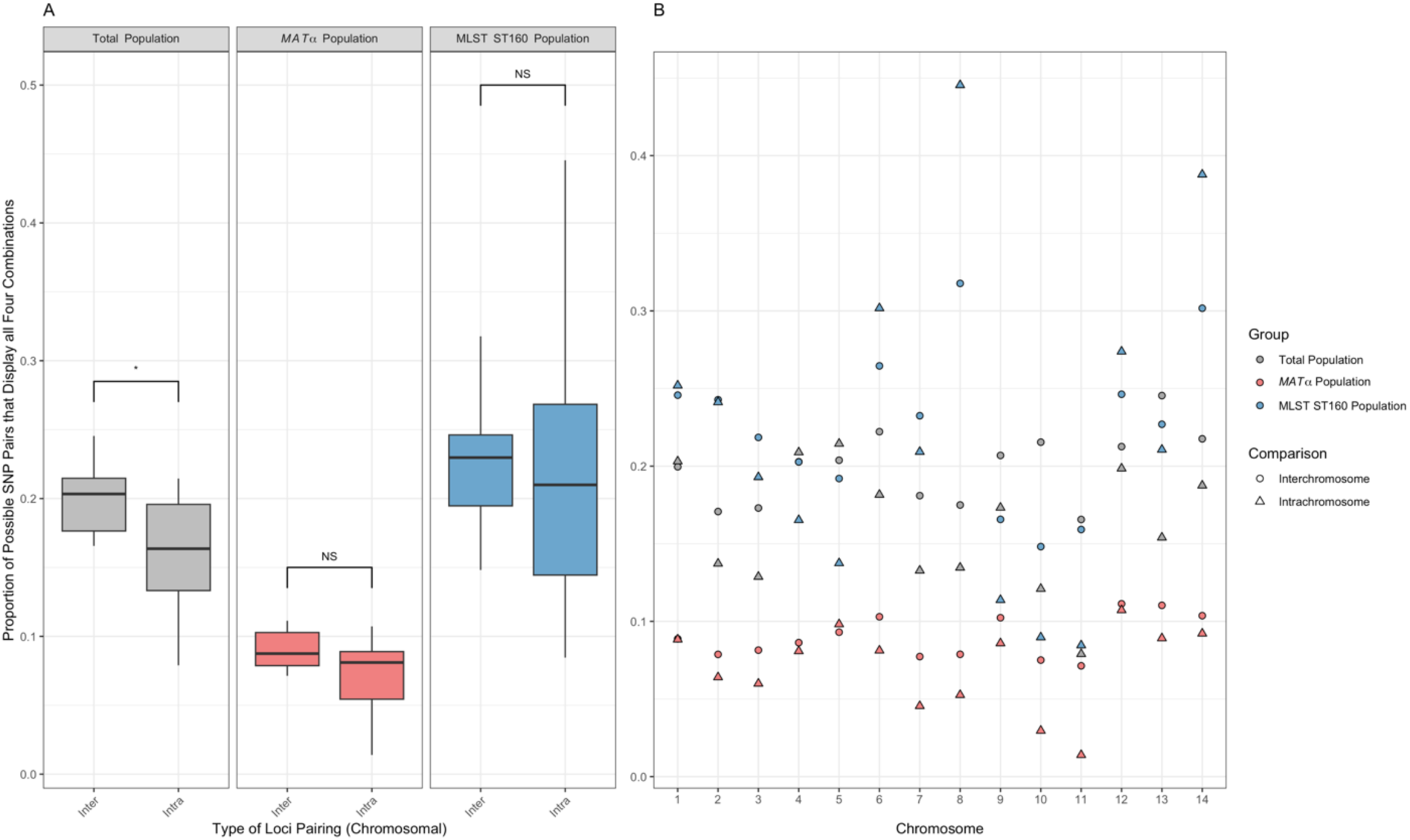
Distribution of the proportions of SNP pairs that display all four genotypes in each population, for both intra-chromosomal SNP pairs and inter-chromosomal SNP pairs. The three different study populations, total population (N = 24), mating type *MAT*α population (N = 23), and MLST ST160 population (N=15) are shown in grey, pink, and blue, respectively. (A) Statistical significance in the difference between inter- and intra-chromosomal SNP pairs being phylogenetically incompatible was measured through the Wilcoxon rank sum exact test. The Boxplot center line represents the median and the top and bottom represent the interquartile range. *p-value < 0.05. (B) Shows the proportion of SNP pairs, per chromosome, that display all four genotypes. Inter-chromosomal and intra-chromosomal SNP pairs are represented by circles and triangles, respectively.

Among the 14 chromosomes in the nuclear genome, there was a range in the proportions of SNP pairs being phylogenetically incompatible in each of the three populations (**Table 3**). In the total population, the lowest proportion was found on chromosome 11 (7.90%) and the highest was found on chromosome 5 (21.45%). In the *MAT*α population, the lowest proportion was found on chromosome 11 (1.39%) and the highest was found on chromosome 12 (10.72%). In the ST160 population, the lowest proportion was found on chromosome 11 (8.46%) and the highest was found on chromosome 8 (44.55%). The proportions of SNP pairs being phylogenetically incompatible across 14 chromosomes in the total population and in the *MAT*α population are highly correlated with each other (Pearson correlation coefficient r = 0.9034, p<0.01). However, neither of them was significantly correlated with the chromosomal proportions in the ST160 population (r < 0.4, p>0.5 in both comparisons).

#### Four-gamete test of mitogenome SNPs

We calculated the mitogenome SNP pairs that are phylogenetically incompatible. In the total population, 36.3% of possible pairs had all four genotypes observed in the population (**Table 3**). This was notably lower in the *MAT*α population where 22.3% displayed all four gametes (**Table 3**). The distributions of SNP pairs showing phylogenetic incompatibilities are shown in **Figure 4**, for both the total population (**Figure 4A**) and the *MAT*α population (**Figure 4B**). We mapped the mitogenome SNPs to the annotation of the JEC21 mitogenome annotation file created by [35]. In both populations, most phylogenetic incompatibilities involved SNPs that were concentrated in a few regions of the mitogenome. These regions were mainly located in non-protein-coding DNA, in-between tRNA genes, and the intronic regions within the *COX1* gene.

**Figure 4.**
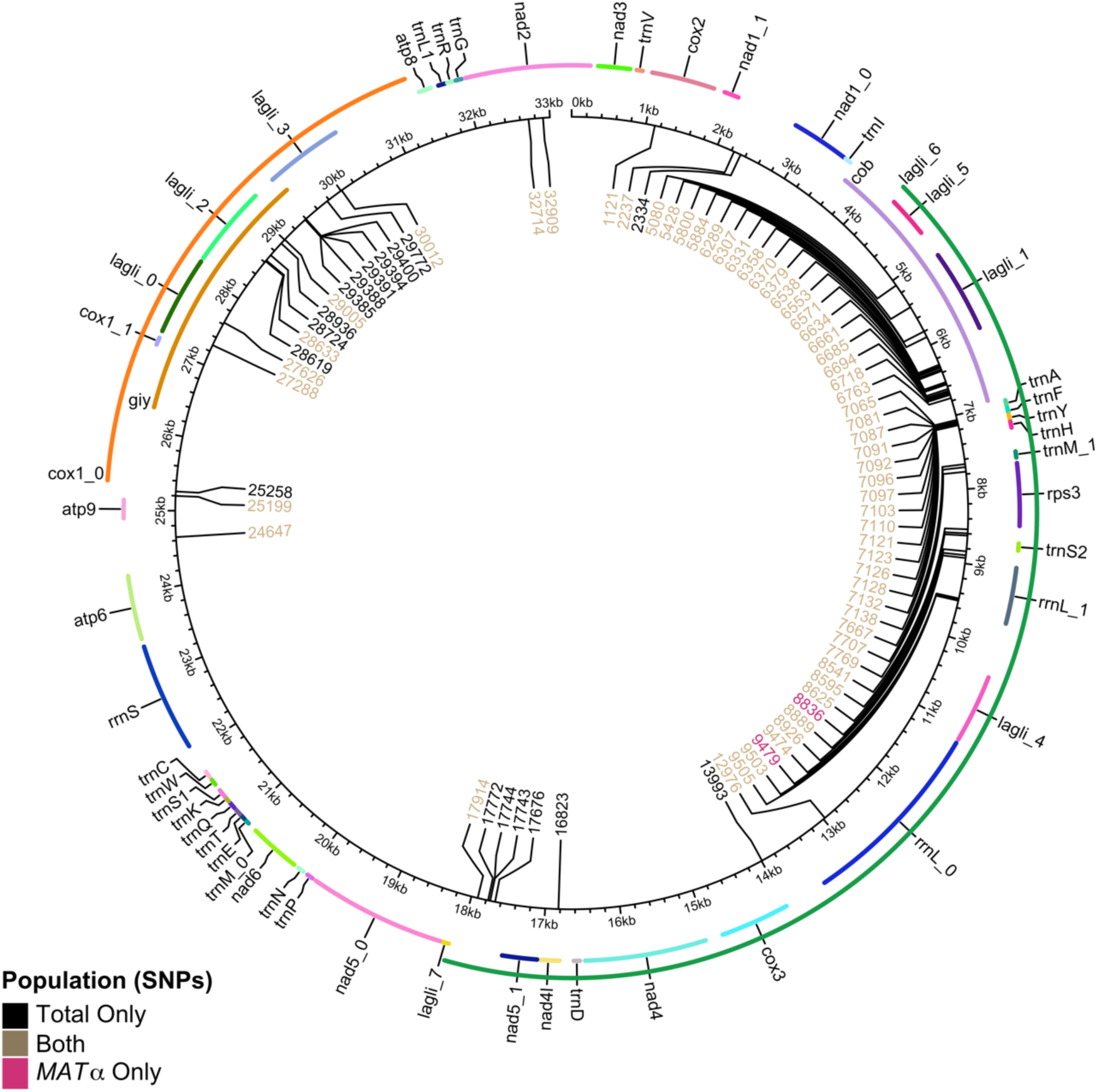
Distribution of biallelic mitochondrial single nucleotide polymorphisms (mtSNPs) across the circular mitochondrial genome (in kilobase pairs). Each mtSNP is denoted by location on the mitogenome in base pairs from an arbitrarily set origin. MtSNPs found only in the total population are coloured black (N=24, 17 loci), only in the *MAT*α population (N=23, 2 loci) in pink, and loci found in both populations in brown (58 loci). Mitochondrial genes are annotated around the perimeter, labeled and coloured only for differentiation purposes.

#### Relationship between physical distance and frequency of phylogenetically incompatible SNP pairs within individual nuclear chromosome and the mitochondrial genome

To assess the distribution of phylogenetically incompatible SNP pairs within chromosomes, we examined the relationship between the SNP pair’s physical distance on each chromosome and the ratio of SNP pairs being phylogenetically incompatible over the total number of pairs of nucleotide sites at each distance. The results are summarized in **Figures 5 and 6**. Across all nuclear chromosomes, a larger number of SNP pairs, normalized by the total number of possible SNP pairs at the same distance within the chromosome, displayed phylogenetic incompatibility as the physical distance between them increased (**Figure 5**). This result is consistent with the observed phylogenetic incompatibility being derived from meiotic crossing-over where SNP pairs located further apart on a chromosome undergo a higher rate of recombination. This analysis was conducted for the total population, the *MAT*α subpopulation, and the ST160 subpopulation, all of which exhibited similar patterns, showing evidence for increased recombination between SNP pairs that are further apart on the same chromosome (**Figure 5**).

**Figure 5.**
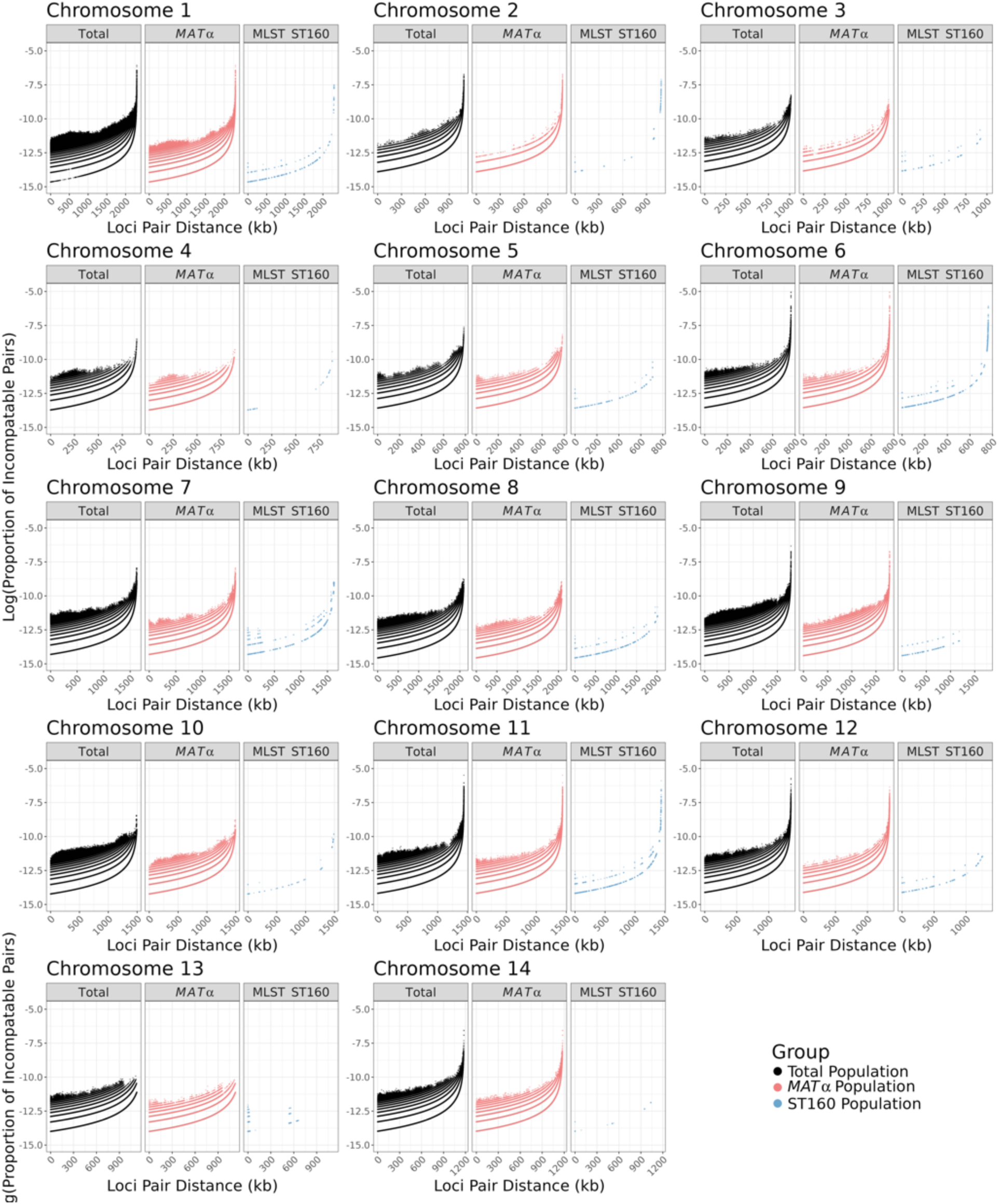
The proportion of biallelic, intrachromosomal SNP pairs exhibiting phylogenetic incompatibility, normalized by the total number of possible SNP pairs at the same distance apart [in kilobases] within the chromosome, displayed on a logarithmic scale. The three different study populations, total population (n = 24), mating type *MAT*α population (n = 23), and MLST ST160 population (n=15) are shown in black, pink, and blue, respectively, for each of the chromosomes. There is a similar trend among the populations, where the proportion of phylogenetically incompatible SNP pairs increases non-linearly as the physical distance between them increases.

**Figure 6.**
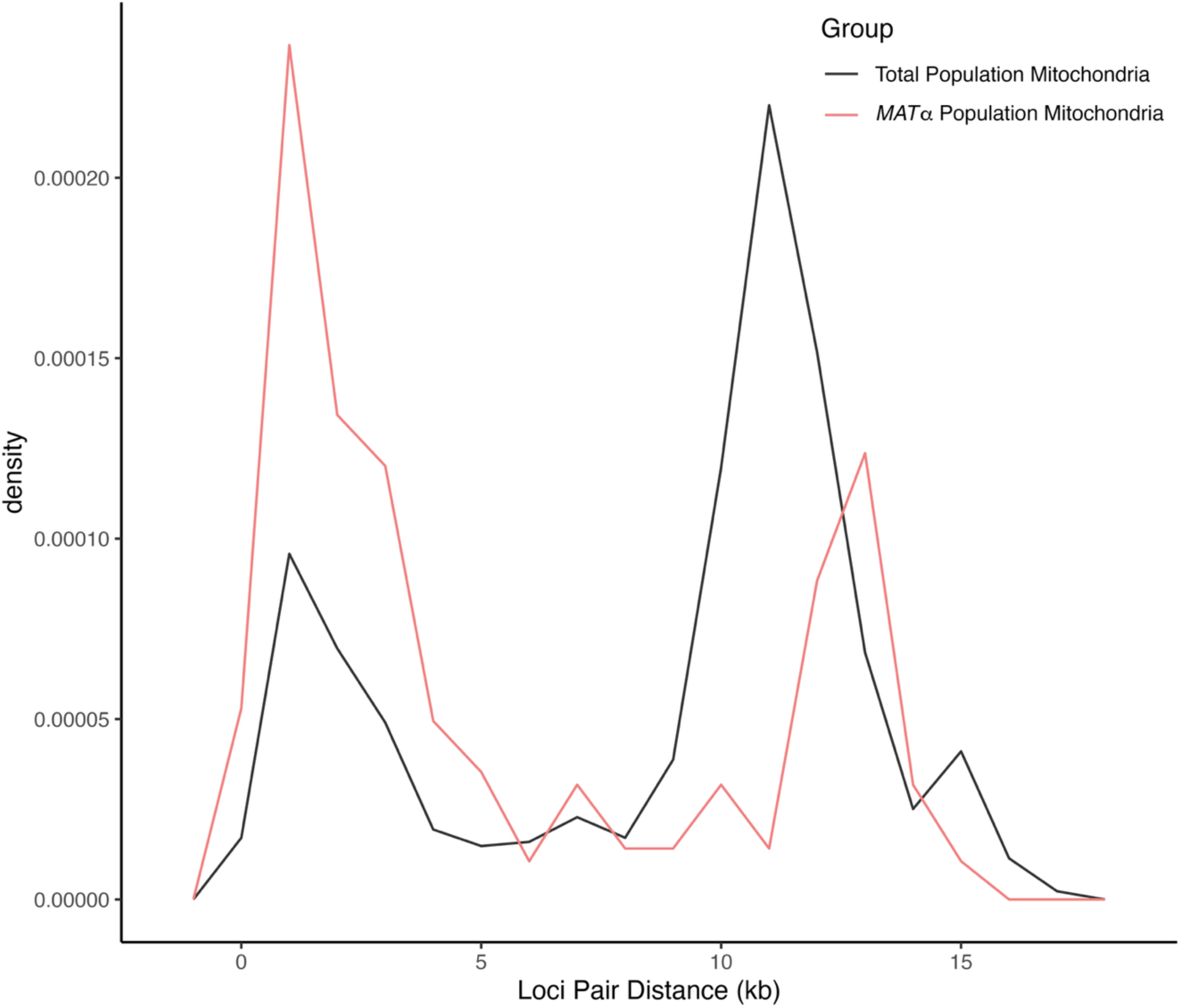
The density of biallelic mitochondrial single nucleotide polymorphisms (mtSNPs) pairs exhibiting phylogenetic incompatibility, calculated based on the shorter distance between each pair (in kilobases) with 1 kb wide bins. The total population (n = 24) and the *MAT*α population (n = 23) are shown in black and pink, respectively. Both populations showed a roughly bimodal distribution of mtSNPs showing all four combinations, with the *MAT*α population showing a higher peak with lower physical distance.

For the mitogenome, both the total population (n=24) and the *MAT*α population (n=23) showed a roughly bimodal distribution of phylogenetic incompatibilities and recombination (**Figure 6**). The results are consistent with uneven distributions and hotspots of recombination events in the mitochondrial genome in this Saudi Arabian population of *C. deneoformans*.

#### Phylogenetic incompatibility between nuclear and mitogenome SNP pairs

To assess the association between nuclear and mitochondrial SNP pairs, we performed a four-gamete test between all pairwise nuclear and mitochondrial SNPs. Here, each nuclear SNP was compared with each mitochondrial SNP in the analyzed populations. In a clonal population without parallel mutation of reversion, we would expect the nuclear and mitochondrial genomes to be completely linked with no evidence of recombination. In contrast, sexual mating would break up the nuclear-mitochondrial SNP associations. Our analyses revealed frequent phylogenetic incompatibilities between nuclear-mitochondrial SNP pairs on most chromosomes (**Figure 7**). Together, these results further support sexual reproduction and recombination in the population.

**Figure 7.**
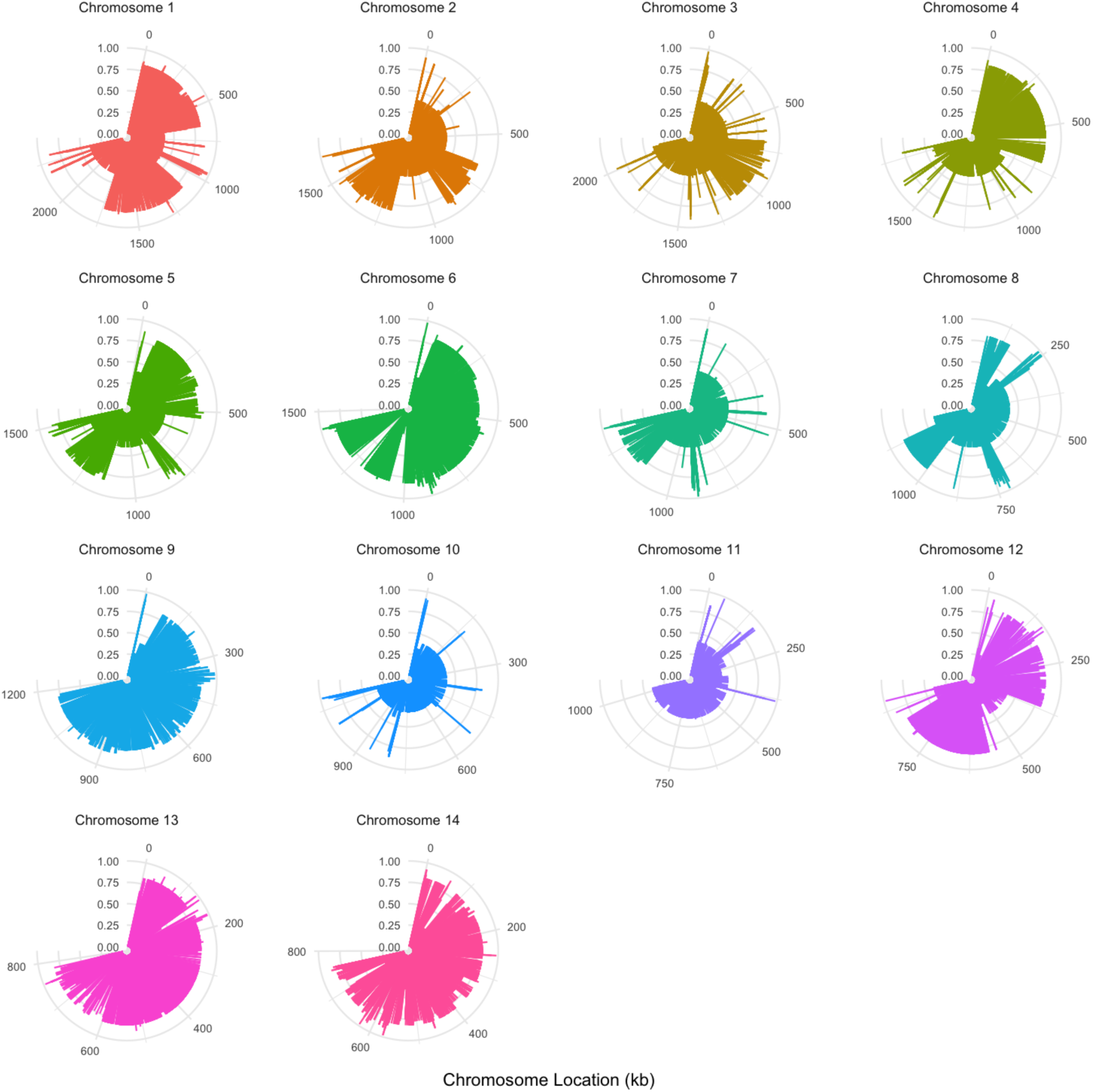
Four gamete test results between nuclear single nucleotide polymorphisms (SNPs) in each chromosome and mitochondrial SNPs (mtSNPs) in the whole population (N=24). The outer circle indicates the position of a chromosomal SNP, in kilobases (kb). Bar height is the proportion of mtSNPs which show phylogenetic incompatibility when paired with SNPs of each specific chromosome. Graphs are scaled such that each chromosome takes up ¾ of a circle, but proportion (radial axis) is consistent for each.

#### Analyses of *MAT*α -locus specific SNPs provide strong evidence of α-α unisexual reproduction

To determine if α-α sexual reproduction contributed to the observed signatures of recombination, we analyzed polymorphisms within the *MAT*α locus (**Figure 8**). Specifically, due to structural differences and sequence divergence between the two mating type loci in *C. deneoformans*, the only type of recombination known to occur within the mating type loci during **a**-α sexual reproduction involves gene conversion [43]. In contrast, α-α unisexual reproduction involving outcrossing between genetically distinct isolates can result in crossing-over within the mating type locus [44]. To investigate whether there is evidence for recombination within the *MAT*α locus, we assessed phylogenetic incompatibility between SNP pairs within the *MAT*α locus using the four-gamete test. Our analysis revealed 10.6% and 3.9% of the SNP pairs located within the *MAT*α locus to be phylogenetically incompatible for the *MAT*α (N=23) population and the ST160 (N=15) population, respectively. Within the ST160 population, the recombination breakpoint that generated the phylogenetic incompatibilities within the mating type locus was between 1612998bp and 1613673bp, a region of 675 bp (**Figure 8**). These results, along with the total SNP counts within the *MAT*α samples, are summarized in **Table 4**. Together, our data provided further evidence for α-α unisexual mating and recombination.

**Figure 8.**
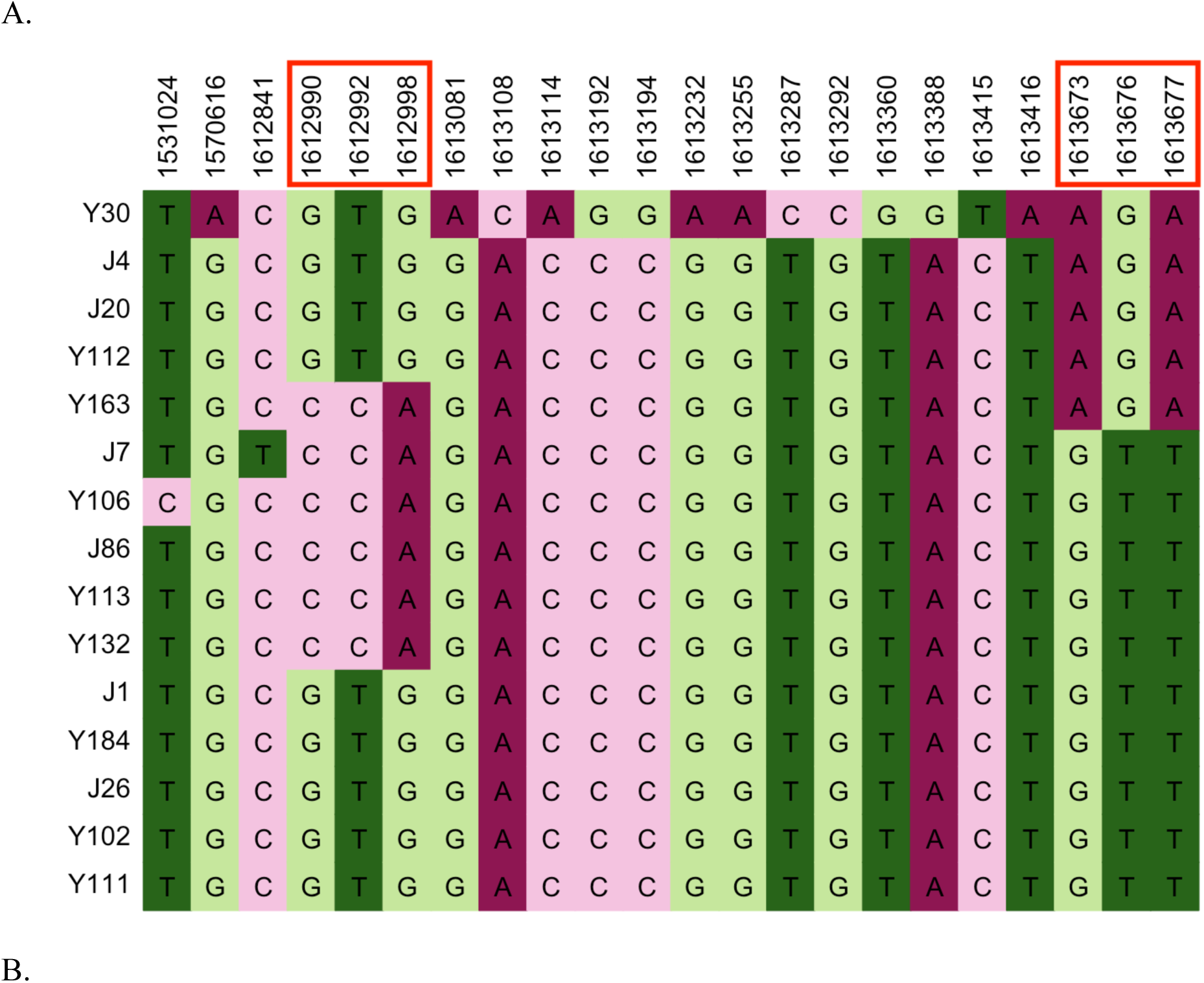
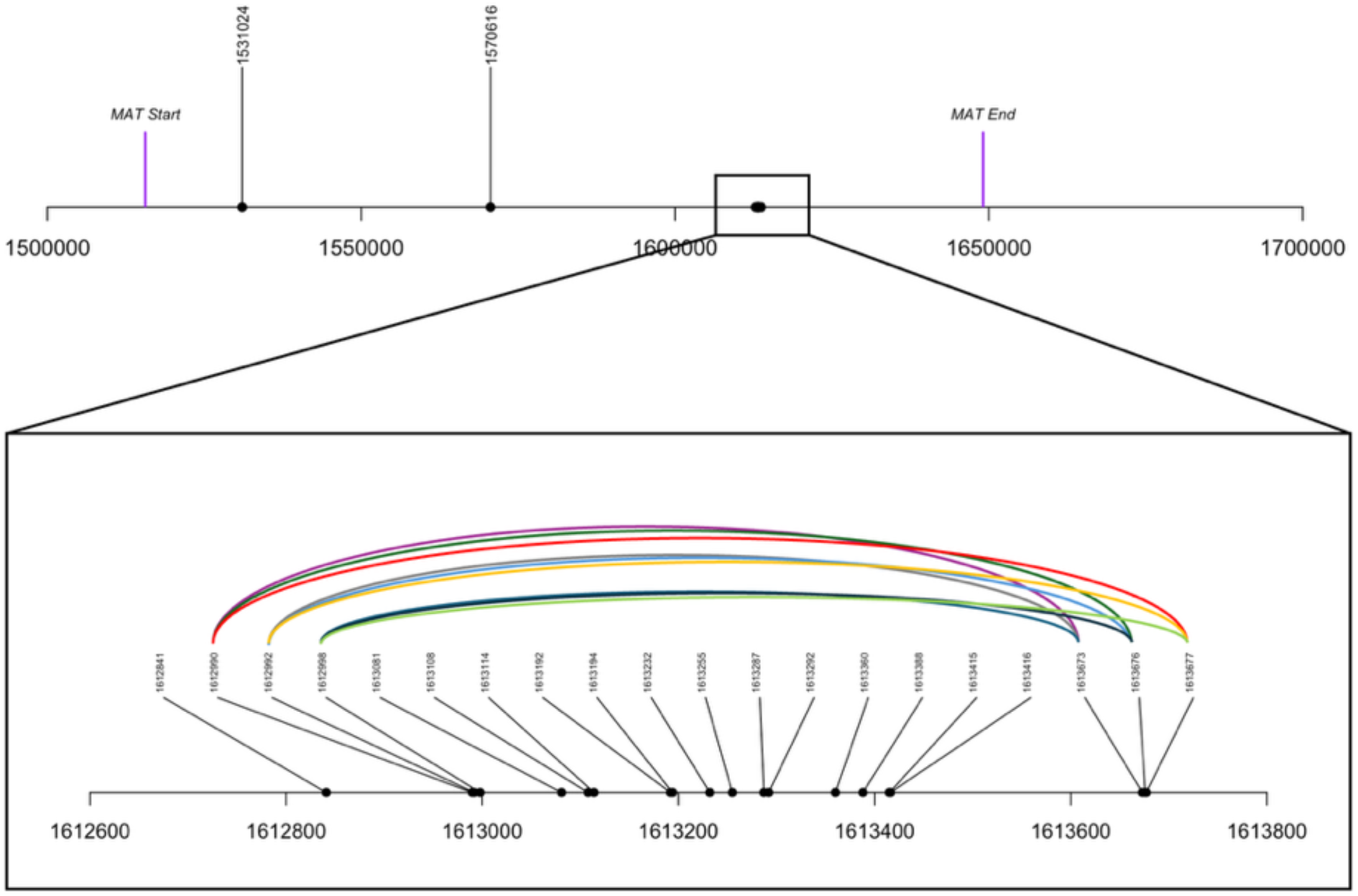
Distribution and association among mitochondrial SNPs located within the *MAT*α locus of 15 ST160 strains of *C. deneoformans* from Saudi Arabia. **A**. Specific nucleotides at the 22 SNP sites. The numbers at the top correspond to the nucleotide positions of the SNPs on Chromosome 4 of the annotated reference genome JEC21. Numbers in red box are the SNPs showing evidence of phylogenetic incompatibility. **B**. Schematic representation of the nine SNP pairs linked by curved colour lines showing evidence of phylogenetic incompatibility.

**Table 4.** Summary of the bi-allelic SNPs located within the *MAT* locus for the *MAT*α and ST160 subpopulations. For each of the two subpopulations, the table includes total SNPs, total number of SNP pair combinations, number of SNP pairs that are phylogenetically incompatible (PI), and the proportion of SNP pairs that are PI. SNP pairs that are phylogenetically incompatible are determined by the four gametes test.

|  | No. of SNPs | No. of pairwise SNP combinations | SNP pairs being PI | PI % |
| --- | --- | --- | --- | --- |
| *MATα* | 531 | 140715 | 14983 | **10.61%** |
| ST160 | 22 | 231 | 9 | **3.90%** |

### Laboratory Mating

Among the 24 isolates crossed with both JEC20 (*MAT***a**) and JEC21 (*MAT*α), 21 successfully mated with JEC20, and consistent with their mating competency and molecular identification of *MAT*α. Similarly, the *MAT***a** isolate Y38 successfully mated with JEC21. Two molecularly identified strains Y97 and Y187 as *MAT*α didn’t show evidence of mating with either JEC20 or JEC21. Analyses of the SNPs in these two strains identified no obvious missense or nonsense mutations in the genes known to be involved in mating in this species (detailed data not shown). None of the 24 isolates showed α-α or **a**-**a** sexual reproduction either on their own or when incubated with either JEC20 or JEC21 in mating type matched co-incubations under our test condition.

## Discussion

In 2019, Samarasinghe et al. discovered an environmental *C. deneoformans* population from the soils along the Red Sea in western Saudi Arabia [28]. This population was identified as the predominant yeast species in the region, revealing a novel ecological niche for this species [28]. Through multi-locus sequence typing (MLST), which analyzes portions of seven housekeeping genes, they concluded the population was largely clonal and had limited evidence for recombination. Interestingly, the population contained mostly MLST genotypes found in other geographic regions as well as a few unique genotypes not yet reported elsewhere [28,29]. This discovery prompted us to further investigate the genomic diversity and mode of reproduction within the population.

In this study, 24 isolates representing the MLST genotypic, geographic, and phenotypic diversities of the original population were sequenced at the whole-genome level. Our analyses revealed abundant SNPs. In total, we identified 104,793 nuclear SNPs and 75 mitochondrial SNPs among the 24 strains, where each SNP had the rare allele present in at least two strains. Notably, 2,027 nuclear SNPs were found among the 15 isolates previously identified as ST160 through MLST [28].

The MLST scheme used in the original genotyping by Samarasinghe et al. identifies isolates based on 3,962 nucleotide positions distributed across seven gene loci located in diverse locations of the nuclear genome [28]. While informative, those seven genes were mostly conserved housekeeping genes. The sequenced portion of those genes together represents 0.0208% (3,962 bp/19,051,922 bp) of the entire genome. Here we identified and analyzed genetic variation among strains in the remaining ∼99.98% of the genomes not covered by the MLST scheme. To determine whether there were inconsistencies in the sequencing results between the original Sanger sequencing conducted for MLST genotyping and the Illumina whole-genome sequencing in this study, we extracted the sequences of the 24 strains at the seven MLST loci from the whole-genome sequence files and compared the sequences in two datasets. Our comparisons found that the sequences of the same strain by the two different methods were all identical to each other. These results indicated that the original MLST sequences and the SNPs identified and analyzed here are robust.

The original analyses based on MLST genotype information indicated several signatures of clonal population structure for the western Saudi Arabian soil population of *C. deneoformans*, including limited nucleotide and allelic diversities, over-representation of several MLST genotypes, and lack of phylogenetic incompatibility between pairs of loci. Here, the identification of abundant SNPs across all chromosomes enabled a detailed analysis of possible recombination. In addition, because the mating types of the strains are known, we analyzed a subset of the strains to identify potential signatures for α-α sexual reproduction among 23 *MAT*α strains. Furthermore, potential signatures of recombination were additionally investigated for a presumed “clone” – 15 strains of ST160. Through these analyses, our results provide robust evidence for both **a**-α and α-α sexual reproduction in nature, along with signals of biparental mitogenome inheritance and recombination in this environmental population.

Initial analysis of the 24 isolates revealed that 19.38% of genome-wide SNP pairs were phylogenetically incompatible, with intra-chromosomal SNP pairs being phylogenetically incompatible ranging from 7.90% to 21.45%, suggesting the presence of frequent recombination. When examining SNP pairs across different distances, a greater proportion of all four gamete combinations was observed as the distance between loci increased. This pattern is consistent with meiotic crossing over and recombination, where SNPs closely located are less likely to have crossing over during meiosis than SNPs located further apart [45,46]. The relatively high homoplasy index of 0.266 also rejects the hypothesis of strict clonal reproduction in this population. These findings align with recent studies indicating that recombination contributes to natural *Cryptococcus* populations more than previously recognized [2,27,47–50].

As an additional test to assess whether recombination was involved in generating the observed genomic diversity, we assessed the associations between nuclear SNPs and mitochondrial SNPs. In a predominantly clonal population, we would see no evidence of phylogenetic incompatibility between nuclear and mitochondrial SNPs. Using the four-gamete test, our analyses identified abundant phylogenetic incompatibilities between nuclear and mitochondrial SNPs, further supporting the hypothesis that that sexual mating occurred in the population.

To estimate the potential contribution of the lone *MAT***a** isolate to the observed phylogenetic incompatibility, we calculated the proportion of phylogenetically compatible SNP pairs among 23 *MAT*α isolates after removing the *MAT***a** isolate. Our analyses revealed that among the 23 *MAT*α isolates, the proportion of phylogenetically incompatible SNP pairs was approximately half (8.76%) of those in the total population (19.38%). The reduction is consistent with **a**-α sexual reproduction playing a significant role in this population.

The high level of recombination detected in the *MAT*α subpopulation could be explained by several cycles of **a**-α sexual reproduction, where *MAT*α alleles are transferred into the *MAT***a** background and then back into *MAT*α through repeated crosses. To further investigate evidence for α-α unisexual reproduction, we measured phylogenetic incompatibility among SNPs within the *MAT* locus. Given the high divergence between **a** and α alleles, homologous recombination within the *MAT* locus is not expected during **a**-α reproduction [51]. However, in α-α sexual reproduction, gene synteny and high sequence similarity enables homologous recombination within the *MAT* locus. Interestingly, our analysis revealed that 10.6% of SNPs within the *MAT*α subpopulation (N=23) were phylogenetically incompatible. In addition, 3.9% of closely located SNPs within the *MAT*α locus were phylogenetically incompatible among the 15 ST160 isolates. These results provide convincing evidence that α-α mating between genetically distinct isolates, crossing-over, and sexual reproduction have occurred in this population.

While the above results suggested α-α mating and sexual reproduction in this population, additional evidence for heterothallic **a**-α mating came from the genome of *MAT*α strain Y87. This strain exhibited a high degree of genetic similarity to the *MAT***a** isolate Y38 in specific regions of the genome, and with discrete regions within the *MAT* locus that alternately aligned with two distinct groups of *MAT***a** isolates. This mosaic pattern suggests historical recombination events involving both **a**-α isolates across the genome and α-α isolates within the *MAT* locus.

Lin et al. were the first to demonstrate that *C. deneoformans* can undergo unisexual reproduction [25,52]. Consistent with our results and the observed mating type distribution, they found that unisexual reproduction among *MAT*α isolates occurs more frequently than between *MAT***a** isolates [25]. Although we did not identify this mutation within our population, recent studies have revealed a naturally occurring loss of function mutation in the GEF Ric8 promotes unisexual reproduction of the lab-generated isolate XL280 that is robustly unisexually fertile [53].

Prior research provides support for α-α unisexual reproduction in natural populations of the human pathogenic *Cryptococcus*. A study of 78 *C. neoformans* serotype A, *MAT*α isolates from India which analyzed MLST loci showed evidence of phylogenetic incompatibility and recombination within the population despite the apparent absence of *MAT***a** mating type in the population [47]. Moreover, population genomic analysis revealed rapid linkage disequilibrium (LD) decay occurring in the VNI serotype A *C. neoformans* global lineage, consistent with recombination in this virtually exclusively *MAT*α lineage in which only four rare *MAT***a** isolates have ever been identified from ∼4000 isolates analyzed (99.9% *MAT*α mating type) [27]. A study in North Carolina identified and characterized three αADα hybrid diploid cells of *C. neoformans* from the environment, and a recent global analysis of the *C. neoformans* AD hybrid population by Cogliati et al. identified additional αADα diploid hybrids [17,52,54]. Similarly, an analysis of serotype A veterinary isolates from Australia revealed αAAα diploids of the serotype A VNII lineage [55]. Additionally, an analysis of multilocus sequence types of the *C. gattii* species complex, the close relative of *CNSC*, suggested that the strains responsible for the ongoing Cryptococcosis outbreak in the Pacific Northwest could have originated from a cryptic α-α unisexual reproduction [56]. Together, these findings demonstrate the importance of α-α unisexual reproduction in natural populations of the human pathogenic *Cryptococcus* [57,58].

It is worth noting that α-α unisexual reproduction in *Cryptococcus* has been linked to biparental mitogenome inheritance in laboratory crosses [58]. Our results here are consistent with α-α unisexual reproduction being associated with biparental mitochondrial inheritance and mitogenome recombination. Mitochondria are essential organelles in eukaryotes, playing critical roles in aerobic respiration, ion homeostasis, cell survival, and virulence in some pathogenic fungi [59,60]. Typically, in **a**-α sexual reproduction, *C. deneoformans* exhibits uniparental mtDNA inheritance, with almost all progeny inheriting the mitogenome from the *MAT***a** parent [23]. However, our analysis of mitochondrial SNPs (mtSNPs) revealed that 36.3% and 22.3% of mtSNP pairs were phylogenetically incompatible within the total sample (N=24) and the *MAT*α subpopulation (N=23), respectively. Both populations exhibit recombination hot spots predominantly localized within a tRNA cluster and non-protein-coding regions of the mitogenome. These regions contain repetitive and palindromic sequences which could increase the likelihood of homologous regions aligning to facilitate recombination without disrupting gene function. This result is also consistent with observations made for recombination hot spots in other yeast species [61]. Mitochondrial recombination was first identified in yeast by Dujon et al. but a mechanism has yet to be defined [62]. Together, these findings add to the growing evidence of mtDNA recombination in natural populations of many fungal species, including the closely related *C. gattii* species complex [35,63–65].

While α-α unisexual reproduction in *C. deneoformans* is known to result in biparental mitochondrial inheritance and mitogenome recombination, phylogenetic incompatibility and recombination in the mitochondrial genome could also be derived from **a**-α sexual reproduction. Previous research by Yan et al. showed that environmental factors, such as elevated temperatures and UV radiation, can influence *C. deneoformans* mitochondrial inheritance, promoting biparental mtDNA inheritance and recombination in **a**-α sexual reproduction [66]. It is possible that environmental conditions in Saudi Arabia may drive biparental mtDNA inheritance and recombination in *C. deneoformans* **a**-α sexual reproduction. However, this would not explain the observed phylogenetic incompatibility observed within the *MAT*α locus. Analyses of genetic crosses involving the Saudi Arabian strains are needed to test this hypothesis further.

Our study provides compelling evidence that the *C. deneoformans* population in Saudi Arabia is far more genetically diverse than previously thought, challenging the assumption that these populations are predominantly clonal. The results here underscore the limitations of MLST in detecting sexual reproduction, suggesting that both **a**-α sexual reproduction and α-α unisexual reproduction are likely more common than previously hypothesized. Further research is needed to explore the ecological and genetic factors driving **a**-α sexual reproduction and α-α unisexual reproduction in nature, mechanisms of mitochondrial recombination, and environmental influences on populations of *CNSC*. Such efforts will be key to better understanding the evolution, virulence, and adaptability of this and other pathogenic fungi.

## Acknowledgements

We thank Paul Thorn for his help with the four-gamete test. This research is supported by Natural Science and Engineering Research Council (NSERC) of Canada (J.X.), by NIH/NIAID R01 grants AI039115-28 and AI050113-20 (J.H.). J.H. is co-director and fellow of the CIFAR program Fungal Kingdom: Threats & Opportunities.

